# Mutagenic effectiveness and efficiency of gamma rays and sodium azide in M_2_ generation of Cowpea [*Vigna unguiculata* (L.) Walp.]

**DOI:** 10.1101/2020.03.09.983486

**Authors:** Aamir Raina, Samiullah Khan

## Abstract

Legumes play a pivotal role in combating the chronic hunger and malnutrition in the developing nations and are also ideal crops to achieve global food and nutrition security. In the era of climate change, erratic rainfalls, depleting arable land and water resource, feeding the rapidly growing population is a challenging task. Like most other pulses, cowpea is a self pollinated, nutritious, versatile and widely adapted grain legume, but harbor a little accessible genetic variability. Lack of sufficient genetic variability and small size of flowers, traditional plant breeding methods are not enough to meet the demands of improvement of cowpea. Hence, induced mutagenesis was employed to induce significant genetic variability across a range of agro-economical traits in two cowpea varieties Gomati VU-89 and Pusa- 578 from M1 to M4 generations. The success of induced mutagenesis largely depends on the selection of appropriate mutagen, its dose, effectiveness and efficiency. Hence present study was conduct to assess the effectiveness and efficiency of single and combined doses of sodium azide and gamma rays to set an appropriate protocol for induced mutagenesis experimentation in cowpea.

## 1. Introduction

Pulses are considered as vital constituents in the diets of people and hence are most suitable crops to achieve the global food and nutrition security [1]. The production of pulses in India is much lower as compared to global pulse production (Table 1). Additional there is a progressive decline in the per capita pulses consumption from 1951 (60.7 g/day) to 2017 (52.9 g/day) [2]. The low yielding potential of crops is one of the primary obstacles in achieving the desired goals of production. The common asynchronous and longer maturity period, more flower drop and less seed set, natural and anthropogenic pressures are other obstacles in achieving the higher production of pulses.

**Table 1.** Average yield of pulses in India and world compared during 2007-2016.

| YEAR | PULSES YEILD (Q/ha) |  |
| --- | --- | --- |
|  | INDIA | WORLD |
| 2007 | 5.970 | 8.154 |
| 2008 | 6.042 | 8.607 |
| 2009 | 6.706 | 9.285 |
| 2010 | 6.496 | 9.043 |
| 2011 | 6.163 | 8.769 |
| 2012 | 6.460 | 9.381 |
| 2013 | 7.162 | 9.745 |
| 2014 | 7.056 | 9.436 |
| 2015 | 6.465 | 9.505 |
| 2016 | 6.644 | 9.929 |

**Source:** FAOSTAT | © FAO Statistics Division, 2017.

Among the ten primary pulse crops as recognised by FAO, Cowpea is an important member on the basis of its use, nutritive value and other desired qualities [3–5]. It is suitable for cultivation in subtropical areas of the world and usually grown alone but can be intercropped with a range of other crops including sorghum, millet, maize, cassava and cotton [6]. Cowpea is considered as a vital constituent of traditional cropping system because of its capability to reinstate soil fertility for succeeding cereal crops grown in rotation with it and enhances soil fertility mostly in smallholder farming systems where very less or no fertilizer is used. Cowpea growing countries in the Asian region are India, Sri Lanka, Bangladesh, Myanmar, China, Korea, Thailand, Indonesia, Nepal, Pakistan, Philippines and Malaysia (Fig 1) [7]. Nigeria is the top most producer of cowpea with an estimated 45% of the world cowpea production (Table 2). The objective of this study was to study effect of mutagens on physiological parameters and their mutagenic effectiveness and efficiency.

**Fig 1.**
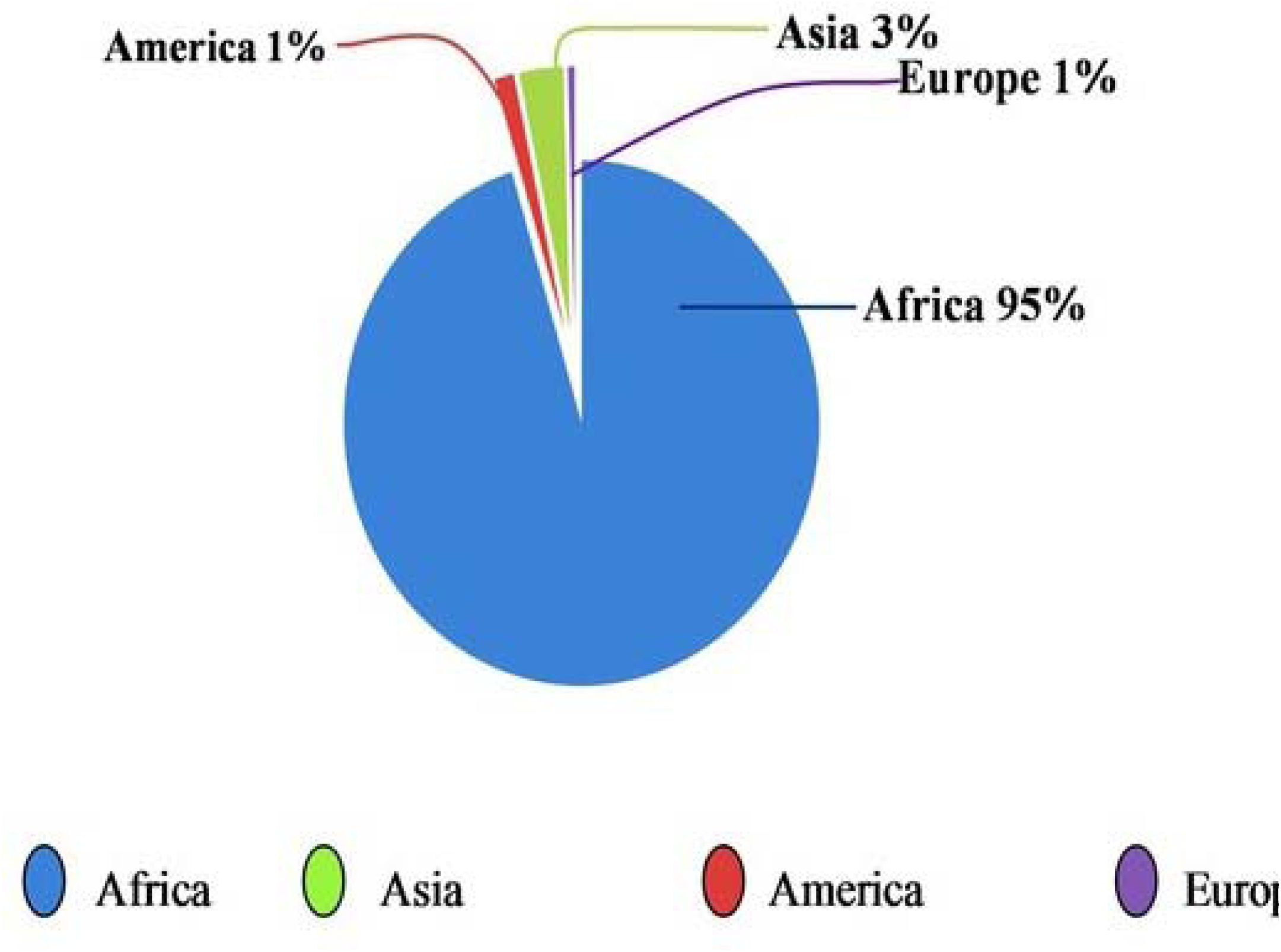
**Cowpea p**roduction.

**Table 2:** World’s top 10 producers of Cowpea (Average 2016).

| **RANK** | **COUNTRY** | **PRODUCTION (mt)** |
| --- | --- | --- |
| 1 | Nigeria | 3.02 |
| 2 | Niger | 1.98 |
| 3 | Burkina Faso | 0.60 |
| 4 | Cameroon | 0.19 |
| 5 | United Republic of Tanzania | 0.18 |
| 6 | Sudan | 0.16 |
| 7 | Mali | 0.14 |
| 8 | Kenya | 0.14 |
| 9 | Myanmar | 0.11 |
| 10 | Sri Lanka | 0.01 |

**Source:** FAOSTAT | © FAO Statistics Division, 2017.

## 2. Materials and methods

### 2.1. Experimental materials and Seed Irradiation

In the present study, M_0_ seeds were obtained from NBPGR, New Delhi (S Table 1). Initially, LD50 was calculated, the doses of gamma rays and sodium azide were optimised by calculating values based on germination rate and survival percentage [6]. A set of 300 dry and healthy M_0_ seeds were pre-soaked in the double distilled water for 9 hours and then treated with 0.01%, 0.02%, 0.03% and 0.04% concentrations of sodium azide (SA) solution for 6 hrs at 25 C at Mutation Breeding Laboratory, Department of Botany, Aligarh Muslim University, Aligarh, India. Another set of 600 M_0_ seeds per dose were irradiated with 100, 200, 300 and 400 Gy of gamma radiation using a Cobalt 60 (^60^Co) gamma source under standard conditions at NBRI, Lucknow, India. Each set of irradiated 300 M_0_ seeds were also treated with respective SA concentration to produce M_1_ seeds. Each set of seeds were washed in running water.

### 2.2. Experimental site and crop cultivation

The experimentation were performed at Agriculture field, Aligarh Muslim University. The net field size included 13 (S Fig 1). Cowpea M_1_ seeds were sown in 10 replications of 30 seeds to raise the M_1_ generation during mid-april 2014-October 2014 along with respective controls. Field was irrigated at regular intervals after seed sowing and; weeding was carried out at seedling and flowering stage. All the M_1_ plants were harvested separately at pod stage to collect M_2_ seeds. During the mid-april 2015-October 2015, 10 healthy M_2_ seeds collected from each M_1_ plant were sown treatment wise for raising M_2_ generation. In M_3_ generation seven and four high yielding stable mutant lines of Gomati VU-89 and Pusa-578, respectively were identified and advanced to M_4_ generation during the mid-april 2017-October 2017 [8].

All the recommended cultivation practices were taken care of; such as 15 - 20 N, 50 - 60 P_2_O_5_ and 50 - 60 K_2_O kilogram per hactare were mixed with soil.

**S1 Fig.** Field plots layout of M_1_ generations.

(Source: Raina et al., 2020. http://creativecommons.org/licenses/by/4.0/).

### 2.3. Field Analysis

#### 2.3.1. Germination of Seeds

Germination of seeds was noted in control and treated sets and the number of seeds germinated in each treated and untreated population was counted. After recording germination counts, the percentage of seed germination was calculated on the basis of total number of seeds sown in the field using the following formulla.

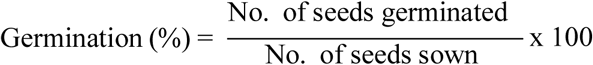

#### 2.3.2. Chlorophyll mutations

These mutations were noted immediately after seed sowing. The identification and categorisation proposed by Gustafsson [9] and Khan [10] was followed using the following formula Sharma [11].

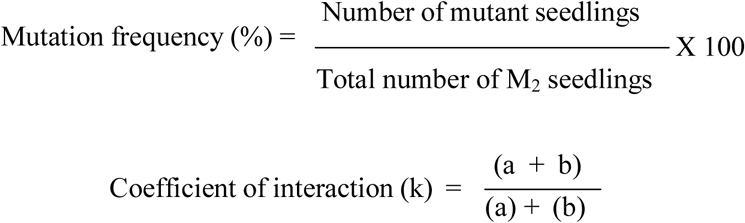

### 2.4. Pollen fertility

The determination of pollen fertility was necessary to assess the pollen sterility based mutagenic effectiveness and efficiency. At the time of flowering, pollens were collected from 30 random plants per treatment and control in M_1_ generation. Pollens were then placed on slides and stained with 1% acetocarmine solution. The uniformly stained pollens were counted as fertile, while as unstained pollens were counted as sterile.

### 2.5. Mutagenic effectiveness and efficiency

The f r e que nc y of m u ta tio n induced by unit mutagen dose refers to mutagenic effectiveness whereas percentage of mutations in relation to biological damage refers to mutagenic efficiency. Mutagenic effectiveness and efficiency was calculated by following formulae given by Konzak *et al*. [12].

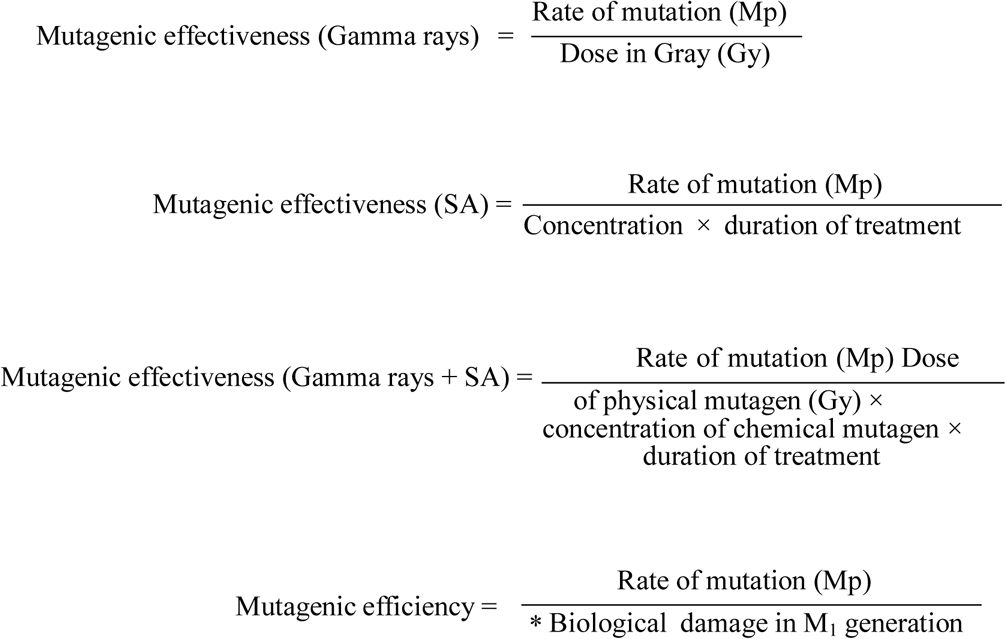

*Biological damage: For measuring the biological damage, three different criteria were used; Injury, Sterility and Meiotic abnormalities refers to %age decrease in seedling height (Mp/I), pollen fertility (Mp/S) and meiotic abnormalities (Mp/Me), respectively.

### 2.6. Morphological mutations

These traits include tall, dwarf, semi dwarf, compact/bushy, prostrate, semi dwarf spreading, one sided branching, axillary branching, broad leaf, narrow leaf, altered leaf architecture, elongated rachis, multiple flower, flower color, open flower, non flowering, late flowering, early maturity,small/narrow pods, bold seeded pods, coat color, coat pattern, shape and surface traits which were examined and modified [13–14]. The characteristics of each mutant is presented in results section. In the field, morphological mutant features for plant height variations, growth habit, leaf, flower, pod and seed were noted.

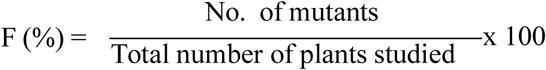

## 3. Results

### 3.1. Seed germination

In M_2_ generation, compared to the control, the gamma irradiated and SA treated population reflected reduced seed germination (Table 3). In both the varieties, germination decreased with the increase in mutagen dose. In the var. Gomati VU-89, control showed a germination percentage of 93.33%, while it reduced from 88.33 to 80.67% in 100 - 400Gy gamma rays and 87.33 to 81.33% in 0.01-0.04% SA and 87.00 to 77.67% in 100Gy + 0.01% SA - 400Gy+0.04% SA treatments. Variety Pusa-578 showed much reduced seed germination, the germination percentage for control was 92.00%, while it decreased from 85.00 to 80.33% in 100 - 400Gy gamma rays and 88.33 to 76.33% in 0.01 - 0.04% SA and 83.67 to 75.33% in 100Gy + 0.01% SA - 400Gy + 0.04% SA in gamma rays+SA treatments. As compared to M_1_ generation, the %age of seed germination was improved in M_2_ generation in both the varieties.

**Table 3.** Effects of gamma rays, SA and their combinations on seed germination and pollen fertility in M_2_ generation two varieties of cowpea

| Treatment | Gomati VU-89 |  |  |  | Pusa-578 |  |  |  |
| --- | --- | --- | --- | --- | --- | --- | --- | --- |
|  | Seed germination | % inhibition | Pollen fertility (%) | % reduction | Seed germination | % inhibition | Pollen fertility (%) | % reduction |
| **Control** | **93.33** | **---** | **96.53** | **---** | **92.00** | **---** | **97.77** | **---** |
| 100Gy $\gamma$ rays | 88.33 | 5.36 | 90.35 | 6.41 | 85.00 | 7.61 | 88.12 | 9.87 |
| 200Gy $\gamma$ rays | 86.00 | 7.86 | 89.11 | 7.69 | 83.33 | 9.42 | 86.88 | 11.14 |
| 300Gy $\gamma$ rays | 82.67 | 11.43 | 85.40 | 11.54 | 81.67 | 11.23 | 81.19 | 16.96 |
| 400Gy $\gamma$ rays | 80.67 | 13.57 | 81.19 | 15.90 | 80.33 | 12.68 | 77.23 | 21.01 |
| **Mean** | **84.42** | **9.55** | **86.51** | **10.38** | **82.58** | **10.24** | **83.35** | **14.75** |
| 0.01% SA | 87.33 | 6.43 | 90.84 | 5.90 | 88.33 | 3.99 | 89.11 | 8.86 |
| 0.02% SA | 86.00 | 7.86 | 87.13 | 9.74 | 84.00 | 8.70 | 84.65 | 13.42 |
| 0.03% SA | 83.33 | 10.71 | 81.68 | 15.38 | 79.67 | 13.41 | 77.72 | 20.51 |
| 0.04% SA | 81.33 | 12.86 | 77.23 | 20.00 | 76.33 | 16.67 | 75.00 | 23.29 |
| **Mean** | **84.50** | **9.46** | **84.22** | **12.76** | **82.17** | **10.69** | **81.62** | **16.52** |
| 100Gy $\gamma$ rays +0.01% SA | 87.00 | 6.79 | 91.58 | 5.13 | 83.67 | 9.06 | 87.13 | 10.89 |
| 200Gy $\gamma$ rays +0.02% SA | 83.67 | 10.36 | 86.14 | 10.77 | 80.67 | 12.32 | 84.41 | 13.67 |
| 300Gy $\gamma$ rays +0.03% SA | 80.33 | 13.93 | 83.66 | 13.33 | 77.33 | 15.94 | 76.49 | 21.77 |
| 400Gy $\gamma$ rays +0.04% SA | 77.67 | 16.79 | 74.50 | 22.82 | 75.33 | 18.12 | 72.03 | 26.33 |
| **Mean** | **82.17** | **11.96** | **83.97** | **13.01** | **79.25** | **13.86** | **80.01** | **18.16** |

### 3.2. Pollen fertility

Pollen fertility was noted in doses employed individually and in combination (Table 3). Its decrease ranged from 6.41% in 100Gy to 15.90% in 400Gy in gamma rays treatments, while as it ranged from 5.90% in 0.01% SA to 20.00% in 0.04% SA treatments and in case of combination treatments it ranged from 5.13% in 100Gy+0.01% SA to 22.82% in 400Gy+0.04% SA. It ranged from 9.87% in 100Gy to 21.01% in 400Gy in gamma rays treatments, while as it ranged from 8.86% to 23.29 in 0.01% SA-0.04% SA in SA treatments and in case of combination treatments it ranged from 10.89% in 100Gy + 0.01% SA to 26.33% in 400Gy + 0.04% SA. The individual mutagen treatments caused more reduction in pollen fertility than combination treatments in both the varieties.

### 3.3. Chlorophyll alterations

Chlorophyll mutations are frequently employed for the determination of mutagenic potency in inducing genetic variability as they are vital indicator in the assessment of induced genetic changes of mutagenized population. A concise account of chlorophyll mutants are presented in Table 4 and Fig 2. The spectrum and frequency of the chlorophyll mutants are presented in Table 5. The chlorophyll mutant frequencies were estimated on M_2_ seedling basis. In both the varieties, except highest dose in each treatment of gamma rays, SA and gamma rays+SA, mutation frequency increased linearly with the increase in mutagen dose. The combination treatment induced higher mutation frequency in comparison to individual treatments. The chlorophyll mutation frequency ranged from 0.72 to 1.06%, 0.88 to 0.92% and 0.53 to 1.68% in gamma rays, SA and gamma rays+SA treatments, respectively. Var. Pusa-578 responded with the lesser mutation frequency compared to the var. Gomati VU-89. In gamma rays, 0.89 to 1.02% in SA treatment and the range of chlorophyll deficient mutants induced by the combined mutagens was 0.48 to 1.78% in the var. Pusa-578.

**Fig 2.**
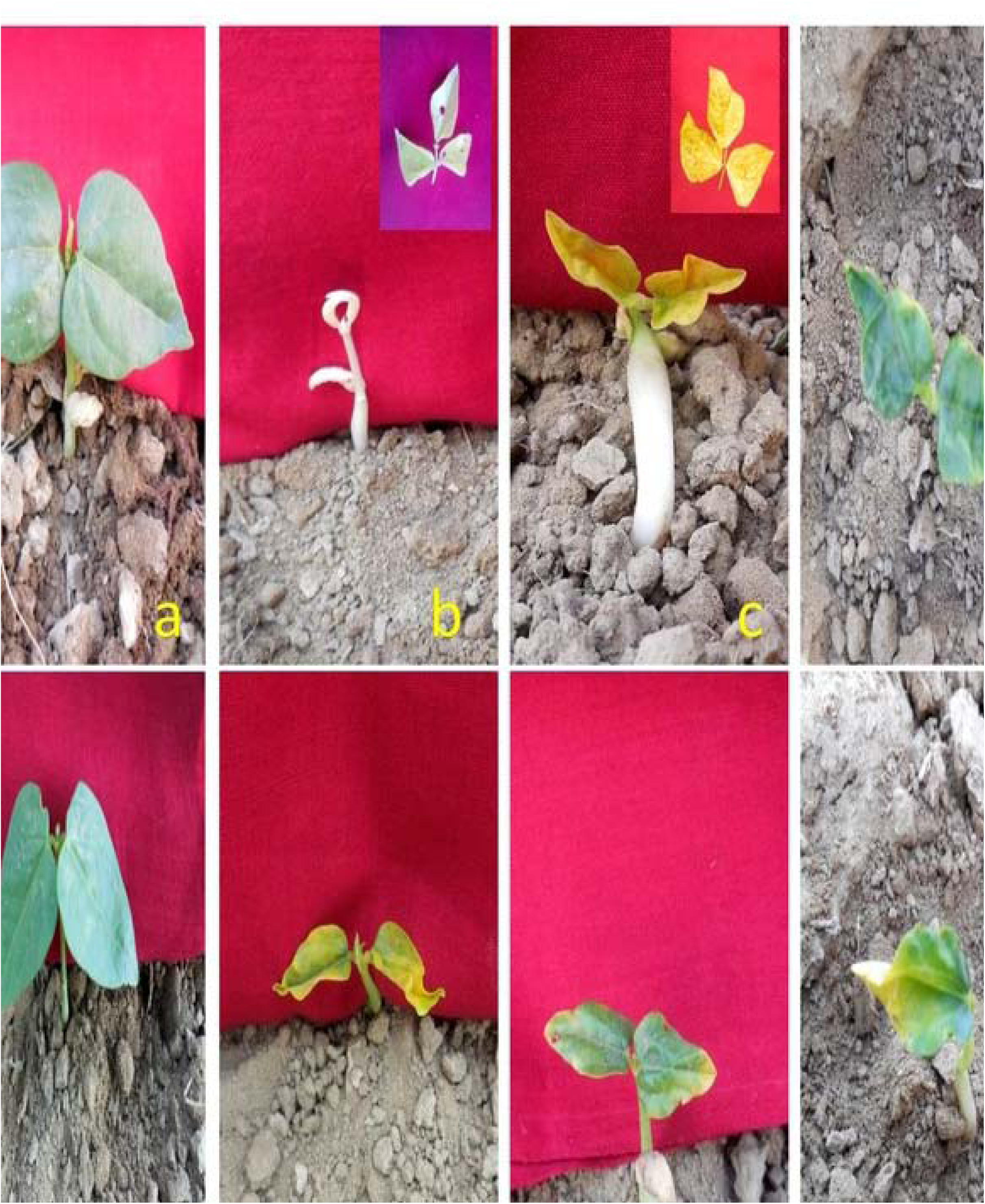
**Chlorophyll mutants.** **a** Gomati VU-89 seedling (Control) showing green color of leaf. **b** Albina **c** Xantha **d** Tigrina **e** Pusa-578 Control **f** Chlorina **g** Xanthaviridis **h** Viridis

**Table 4.**
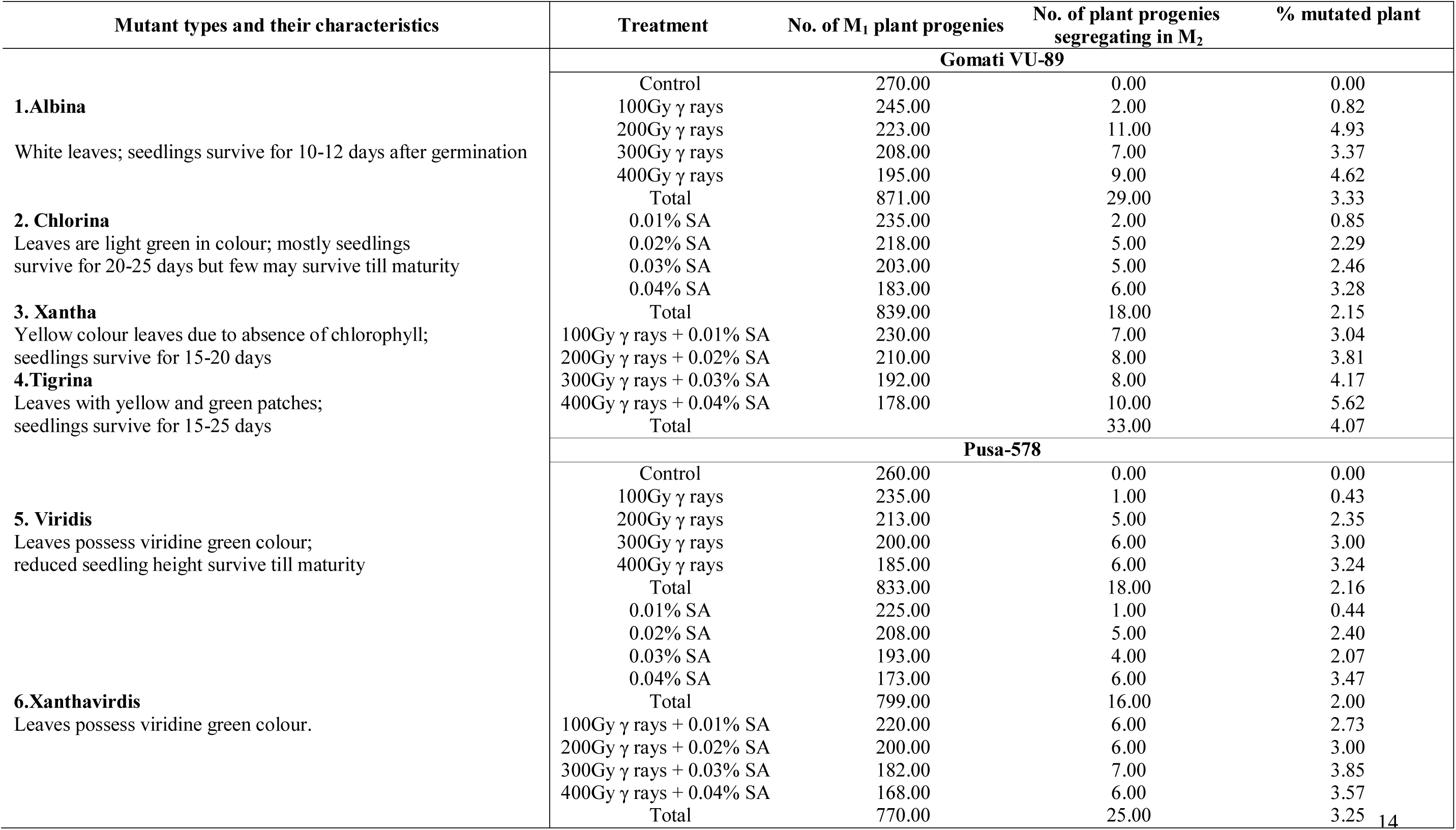
Characteristic cowpea chlorophyll mutants

**Table 5.**
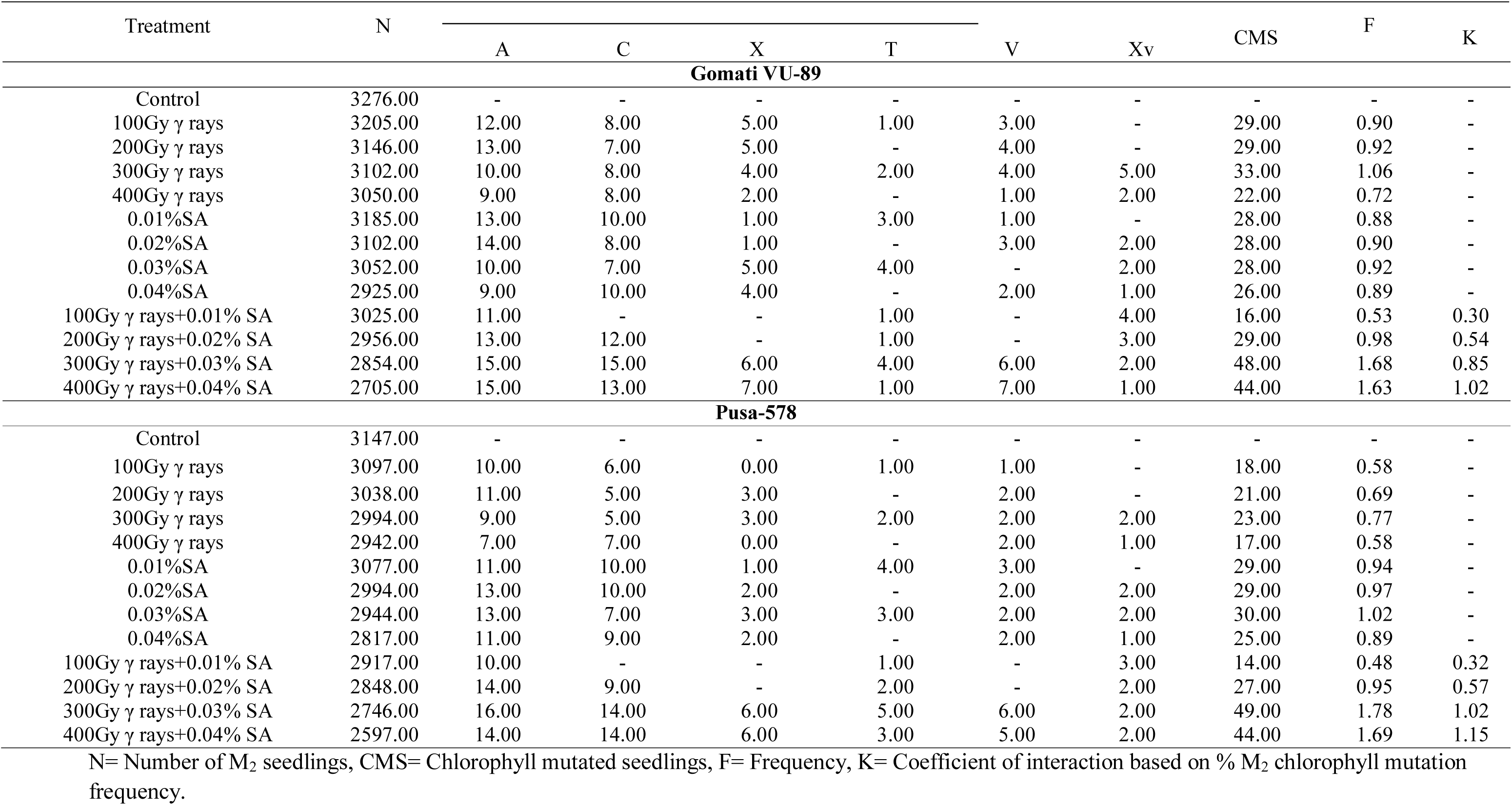
Frequency of chlorophyll mutants.

| Treatment | N |  |  |  |  | V | Xv | CMS | F | K |  |
| --- | --- | --- | --- | --- | --- | --- | --- | --- | --- | --- | --- |
|  |  | A | C | X | T |  |  |  |  |  |  |
| Gomati VU-89 |  |  |  |  |  |  |  |  |  |  |  |
| Control | 3276.00 | - | - | - | - | - | - | - | - | - |  |
| 100Gy $\gamma$ rays | 3205.00 | 12.00 | 8.00 | 5.00 | 1.00 | 3.00 | - | 29.00 | 0.90 | - | |
| 200Gy $\gamma$ rays | 3146.00 | 13.00 | 7.00 | 5.00 | - | 4.00 | - | 29.00 | 0.92 | - | |
| 300Gy $\gamma$ rays | 3102.00 | 10.00 | 8.00 | 4.00 | 2.00 | 4.00 | 5.00 | 33.00 | 1.06 | - | |
| 400Gy $\gamma$ rays | 3050.00 | 9.00 | 8.00 | 2.00 | - | 1.00 | 2.00 | 22.00 | 0.72 | - | |
| 0.01%SA | 3185.00 | 13.00 | 10.00 | 1.00 | 3.00 | 1.00 | - | 28.00 | 0.88 | - |  |
| 0.02%SA | 3102.00 | 14.00 | 8.00 | 1.00 | - | 3.00 | 2.00 | 28.00 | 0.90 | - |  |
| 0.03%SA | 3052.00 | 10.00 | 7.00 | 5.00 | 4.00 | - | 2.00 | 28.00 | 0.92 | - |  |
| 0.04%SA | 2925.00 | 9.00 | 10.00 | 4.00 | - | 2.00 | 1.00 | 26.00 | 0.89 | - |  |
| 100Gy $\gamma$ rays+0.01% SA | 3025.00 | 11.00 | - | - | 1.00 | - | 4.00 | 16.00 | 0.53 | 0.30 | |
| 200Gy $\gamma$ rays+0.02% SA | 2956.00 | 13.00 | 12.00 | - | 1.00 | - | 3.00 | 29.00 | 0.98 | 0.54 | |
| 300Gy $\gamma$ rays+0.03% SA | 2854.00 | 15.00 | 15.00 | 6.00 | 4.00 | 6.00 | 2.00 | 48.00 | 1.68 | 0.85 | |
| 400Gy $\gamma$ rays+0.04% SA | 2705.00 | 15.00 | 13.00 | 7.00 | 1.00 | 7.00 | 1.00 | 44.00 | 1.63 | 1.02 | |
| Pusa-578 |  |  |  |  |  |  |  |  |  |  |  |
| Control | 3147.00 | - | - | - | - | - | - | - | - | - |  |
| 100Gy $\gamma$ rays | 3097.00 | 10.00 | 6.00 | 0.00 | 1.00 | 1.00 | - | 18.00 | 0.58 | - | |
| 200Gy $\gamma$ rays | 3038.00 | 11.00 | 5.00 | 3.00 | - | 2.00 | - | 21.00 | 0.69 | - | |
| 300Gy $\gamma$ rays | 2994.00 | 9.00 | 5.00 | 3.00 | 2.00 | 2.00 | 2.00 | 23.00 | 0.77 | - | |
| 400Gy $\gamma$ rays | 2942.00 | 7.00 | 7.00 | 0.00 | - | 2.00 | 1.00 | 17.00 | 0.58 | - | |
| 0.01%SA | 3077.00 | 11.00 | 10.00 | 1.00 | 4.00 | 3.00 | - | 29.00 | 0.94 | - |  |
| 0.02%SA | 2994.00 | 13.00 | 10.00 | 2.00 | - | 2.00 | 2.00 | 29.00 | 0.97 | - |  |
| 0.03%SA | 2944.00 | 13.00 | 7.00 | 3.00 | 3.00 | 2.00 | 2.00 | 30.00 | 1.02 | - |  |
| 0.04%SA | 2817.00 | 11.00 | 9.00 | 2.00 | - | 2.00 | 1.00 | 25.00 | 0.89 | - |  |
| 100Gy $\gamma$ rays+0.01% SA | 2917.00 | 10.00 | - | - | 1.00 | - | 3.00 | 14.00 | 0.48 | 0.32 | |
| 200Gy $\gamma$ rays+0.02% SA | 2848.00 | 14.00 | 9.00 | - | 2.00 | - | 2.00 | 27.00 | 0.95 | 0.57 | |
| 300Gy $\gamma$ rays+0.03% SA | 2746.00 | 16.00 | 14.00 | 6.00 | 5.00 | 6.00 | 2.00 | 49.00 | 1.78 | 1.02 | |
| 400Gy $\gamma$ rays+0.04% SA | 2597.00 | 14.00 | 14.00 | 6.00 | 3.00 | 5.00 | 2.00 | 44.00 | 1.69 | 1.15 | |

N= Number of M<sub>2</sub> seedlings, CMS= Chlorophyll mutated seedlings, F= Frequency, K= Coefficient of interaction based on % M<sub>2</sub> chlorophyll mutation frequency.

The spectrum of chlorophyll mutations is given in (Table 6). Among the mutants the occurrence of albina type of chlorophyll mutants was the highest in varieties Gomati VU-89 and Pusa-578. Among the doses the frequency of albina mutants was highest in individual and combination doses of gamma rays+SA, whereas SA treatments induced higher frequency of chlorina mutants in varieties Gomati VU-89 and Pusa-578 (Fig 3).

**Fig 3.**
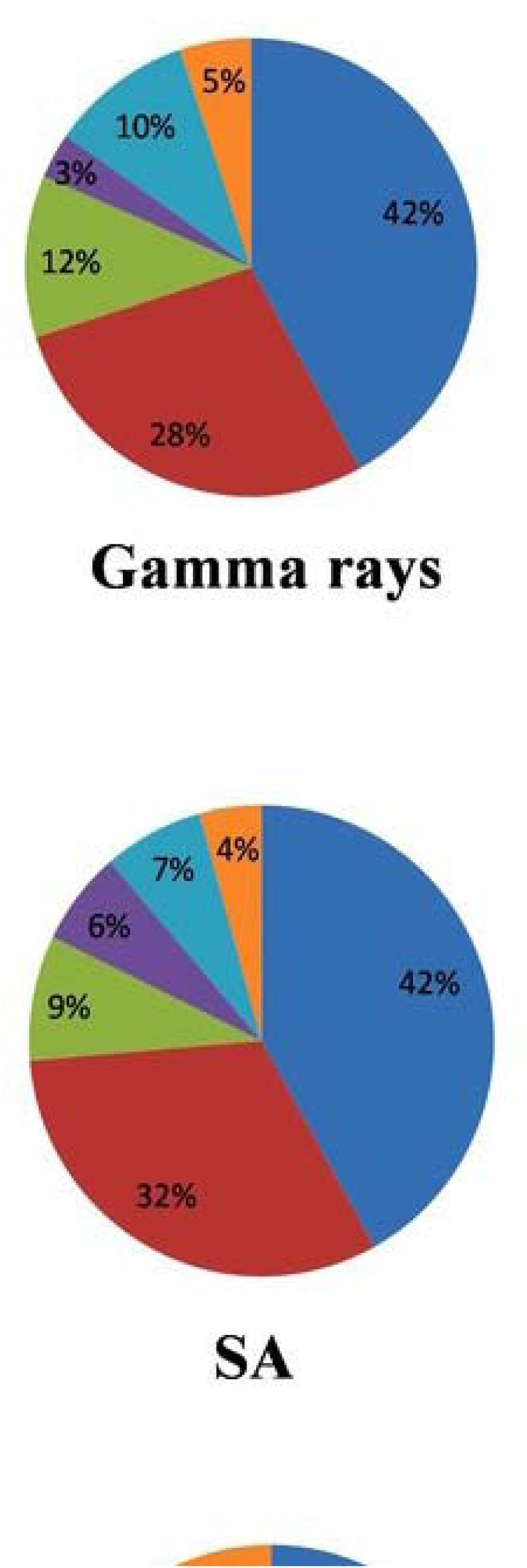
**F**requency of chlorophyll mutations.

**Table 6.**
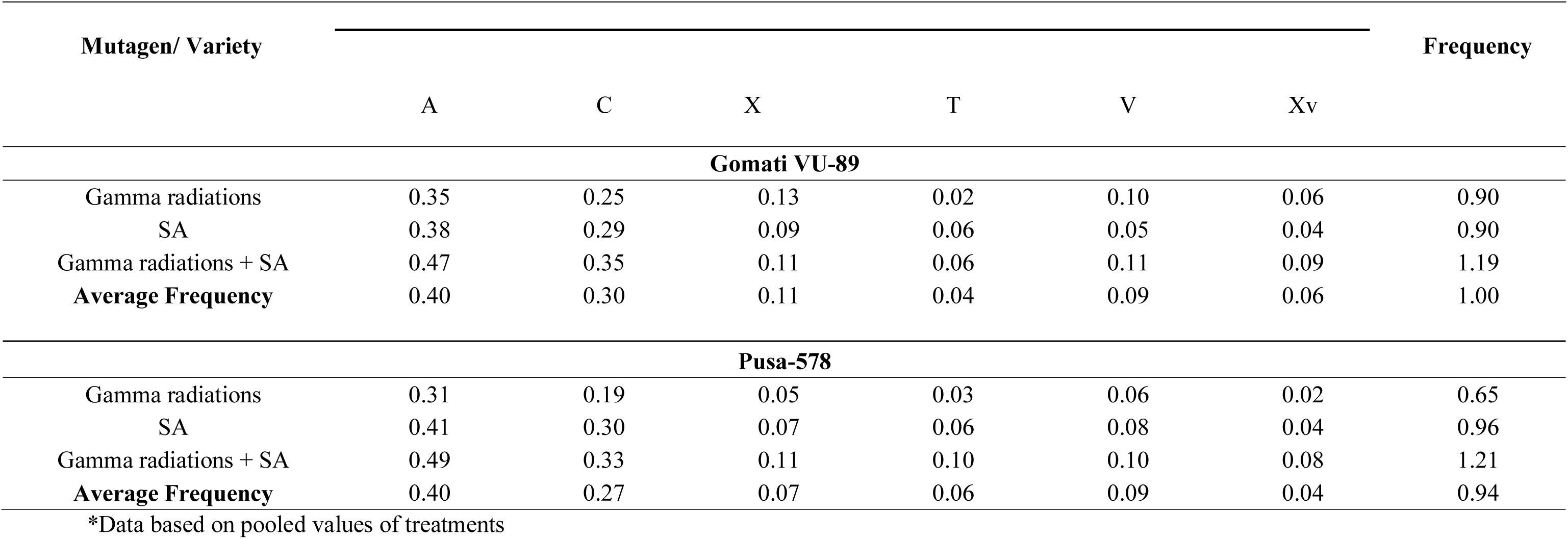
**C**hlorophyll mutation frequency.

### 3.4. Mutagenic effectiveness and efficiency

The mutagen effectiveness and efficiency in a induced mutagenesis determines its usefulness and applicability for crop improvement. Its assessment has been evaluated on the basis of chlorophyll mutation frequency (Tables 7 and 8). The effectiveness and efficiency were highest at the intermediate treatments of mutagens, however lower doses of combined mutagen treatments were most effective and efficient in both the varieties. SA was effective than gamma rays. In both the varieties, effectiveness of single and combined mutagens treatments decreased progressively with the increase in dose in both the varieties. The effectiveness of SA treatments ranged from 25.00 to 41.66 and 16.66 to 41.66 in the varieties Gomati VU-89 and Pusa-578, respectively.

**Table 7.** Effectiveness and efficiency of gamma rays, SA and their combinations in cowpea var. Gomati VU-89.

| Treatment | (%)<br>Seedling<br>injury<br>(I) | (%)<br>Pollen<br>sterility<br>(S) | % | % | Mutagenic<br>effectiveness | Mutagenic efficiency |  |  |
| --- | --- | --- | --- | --- | --- | --- | --- | --- |
|  |  |  | meiotic<br>abnormalities<br>(Me) | Mutated<br>plant<br>progenies<br>(Mp) |  | Mp/I | Mp/S | Mp/Me |
| 100Gy $\gamma$ rays | 10.91 | 6.41 | 3.19 | 2.00 | 0.02 | 0.18 | 0.31 | 0.63 |
| 200Gy $\gamma$ rays | 16.37 | 7.69 | 5.14 | 11.00 | 0.06 | 0.67 | 1.43 | 2.14 |
| 300Gy $\gamma$ rays | 18.88 | 11.54 | 5.47 | 7.00 | 0.02 | 0.37 | 0.61 | 1.28 |
| 400Gy $\gamma$ rays | 25.61 | 15.90 | 8.51 | 9.00 | 0.02 | 0.35 | 0.57 | 1.06 |
| 0.01%SA | 17.40 | 5.90 | 1.63 | 2.00 | 33.33 | 0.11 | 0.34 | 1.23 |
| 0.02%SA | 19.64 | 9.74 | 3.50 | 5.00 | 41.66 | 0.25 | 0.51 | 1.43 |
| 0.03%SA | 22.31 | 15.38 | 6.02 | 5.00 | 27.77 | 0.22 | 0.33 | 0.83 |
| 0.04%SA | 29.13 | 20.00 | 7.63 | 6.00 | 25.00 | 0.21 | 0.30 | 0.79 |
| 100Gy $\gamma$ rays+ 0.01% SA | 16.89 | 5.13 | 5.44 | 7.00 | 1.16 | 0.41 | 1.37 | 1.29 |
| 200Gy $\gamma$ rays +0.02% SA | 20.35 | 10.77 | 6.43 | 8.00 | 0.33 | 0.39 | 0.74 | 1.25 |
| 300Gy $\gamma$ rays+0.03% SA | 25.05 | 13.33 | 9.05 | 8.00 | 0.14 | 0.32 | 0.60 | 0.88 |
| 400Gy $\gamma$ rays+0.04% SA | 31.79 | 22.82 | 10.31 | 10.00 | 0.10 | 0.31 | 0.44 | 0.97 |

**Table 8.** Effectiveness and efficiency of gamma rays, SA and their combinations in cowpea var. Pusa-578.

| Treatments | (% )<br>Seedling<br>injury | (%)<br>Pollen<br>injury | %<br>abnormalities | %<br>plant<br>progenies | Mutagenic<br>effectiveness | Mutagenic efficiency |  |  |
| --- | --- | --- | --- | --- | --- | --- | --- | --- |
|  | (I) | (S) | (Me) | (Mp) |  | Mp/I | Mp/S | Mp/Me |
| 100Gy $\gamma$ rays | 12.28 | 9.87 | 3.19 | 1.00 | 0.01 | 0.08 | 0.10 | 0.31 |
| 200Gy $\gamma$ rays | 17.17 | 11.14 | 5.77 | 6.00 | 0.03 | 0.35 | 0.54 | 1.04 |
| 300Gy $\gamma$ rays | 21.51 | 16.96 | 5.83 | 6.00 | 0.02 | 0.28 | 0.35 | 1.03 |
| 400Gy $\gamma$ rays | 23.47 | 21.01 | 8.48 | 6.00 | 0.02 | 0.26 | 0.29 | 0.71 |
| 0.01%SA | 11.99 | 8.86 | 1.05 | 1.00 | 16.66 | 0.08 | 0.11 | 0.95 |
| 0.02%SA | 14.84 | 13.42 | 3.56 | 5.00 | 41.66 | 0.34 | 0.37 | 1.41 |
| 0.03%SA | 19.18 | 20.51 | 5.00 | 4.00 | 22.22 | 0.21 | 0.20 | 0.80 |
| 0.04%SA | 20.12 | 23.29 | 6.62 | 6.00 | 25.00 | 0.30 | 0.26 | 0.91 |
| 100Gy $\gamma$ rays+ 0.01% SA | 13.11 | 10.89 | 4.05 | 5.00 | 1.00 | 0.38 | 0.46 | 1.23 |
| 200Gy $\gamma$ rays+0.02% SA | 17.72 | 13.67 | 4.90 | 6.00 | 0.25 | 0.34 | 0.44 | 1.22 |
| 300Gy $\gamma$ rays+0.03% SA | 21.12 | 21.77 | 5.63 | 7.00 | 0.13 | 0.33 | 0.32 | 1.24 |
| 400Gy $\gamma$ rays+0.04% SA | 26.88 | 26.33 | 8.43 | 8.00 | 0.06 | 0.30 | 0.30 | 0.95 |

The highest efficiency was noted at 200 Gy of gamma rays followed by 100 Gy+0.01% SA and 0.02% SA in both the varieties of cowpea. Mp/Me based efficiency was maximum at 200Gy gamma rays followed by 0.02% SA and 100Gy+0.01% SA in the var. Gomati VU-89. However, the trend was slightly altered in var. Pusa-578 as the highest efficiency was achieved in 0.02% SA than 100Gy+ 0.01% SA and 100 Gy gamma rays. In both the varieties injury and sterility based mutagenic efficiency was found in the decreasing order i.e., gamma rays> gamma rays+ SA> SA. In var. Gomati VU-89 and Pusa-578, meiotic abnormalities based efficiency was higher in gamma rays + SA treatments gamma rays treatments and SA. Meiotic abnormalities based mutagenic efficiency was higher than injury and sterility based efficiency.

### 3.5. Morphological mutations

A diverse range of mutagen induced morphological mutants were obtained in the mutagenized cowpea population in M2 generation, many of which may be useful from yield improvement prospectives. These mutants revealed differences in the traits such as plant height, growth habit, leaf, flower pod and seed. The morphological mutations, detected at different growth stages of M2 plants, were categorically inspected throughout the growing season. The maximum frequency was recorded in gamma rays+SA treatments (5.86 and 6.56%) and the minimum frequency with gamma rays treatments (4.49 and 5.05%), while the SA treatments induced intermediate frequency of morphological mutants (5.38 and 6.05%) in the varieties (Tables 9 and 10). The overall frequencies of morphological mutants were more in Pusa-578 than Gomati VU-89. The spectrum of morphological mutations is given in (Fig 4). The most prominent morphological mutations were found to be associated with the seed and flower and growth habit, plant size and pod length (Tables 11 and 12). The collective treatments produced significant effects on morphological mutations.

**Fig 4.**
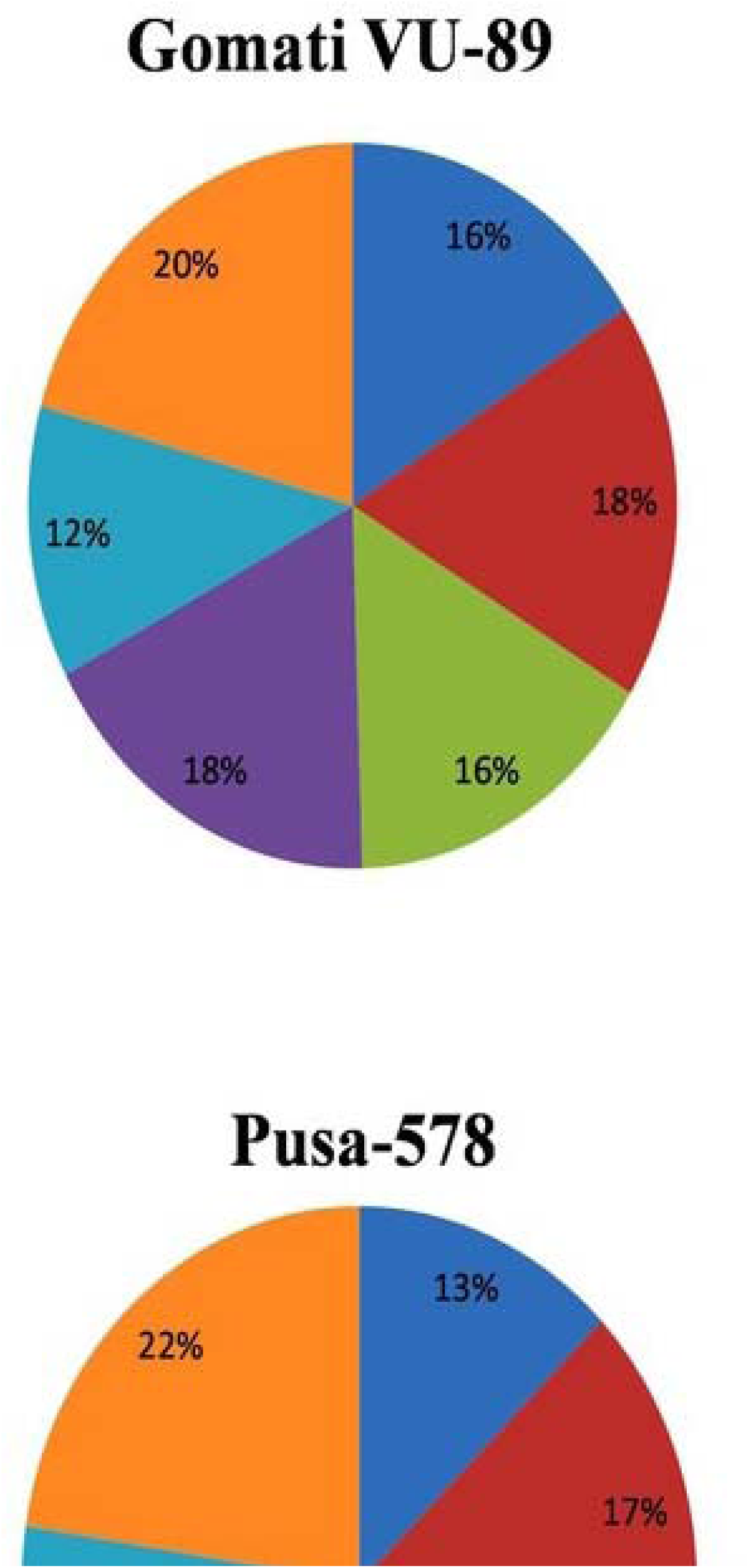
**P**ercentage and spectrum of morphological mutations

**Table 9.** Frequency and spectrum of morphological mutants.

| T | N | Morphological mutant types |  |  |  |  |  | TMP | % Mutated | k |
| --- | --- | --- | --- | --- | --- | --- | --- | --- | --- | --- |
|  |  | H | G | L | F | P | S |  |  |  |
| Control | 3155.00 | - | - | - | - | - | - | - | - | - |
| 100Gy $\gamma$ rays | 3000.00 | 8.00 | - | 6.00 | 2.00 | 9.00 | 4.00 | 29.00 | 0.97 | - |
| 200Gy $\gamma$ rays | 2941.00 | 4.00 | 10.00 | 13.00 | 5.00 | - | 5.00 | 37.00 | 1.26 | - |
| 300Gy $\gamma$ rays | 2897.00 | - | 16.00 | 9.00 | 7.00 | - | 6.00 | 38.00 | 1.31 | - |
| 400Gy $\gamma$ rays | 2845.00 | 8.00 | 8.00 | - | 2.00 | 6.00 | 3.00 | 27.00 | 0.95 | - |
| **Total** | **11683.00** | **20.00** | **34.00** | **28.00** | **16.00** | **15.00** | **18.00** | **131.00** | **4.49** | **0.00** |
| 0.01%SA | 2980.00 | 9.00 | - | 11.00 | 12.00 | - | 3.00 | 35.00 | 1.17 | - |
| 0.02%SA | 2897.00 | 5.00 | - | 6.00 | 5.00 | 11.00 | 6.00 | 33.00 | 1.14 | - |
| 0.03%SA | 2847.00 | - | 12.00 | 7.00 | 10.00 | - | 7.00 | 36.00 | 1.26 | - |
| 0.04%SA | 2720.00 | 10.00 | 7.00 | 13.00 | 8.00 | 7.00 | 4.00 | 49.00 | 1.80 | - |
| **Total** | **11444.00** | **24.00** | **19.00** | **37.00** | **35.00** | **18.00** | **20.00** | **153.00** | **5.38** | **0.00** |
| 100Gy $\gamma$ rays +0.01% SA | 2820.00 | 6.00 | - | - | 15.00 | - | 9.00 | 30.00 | 1.06 | 0.44 |
| 200Gy $\gamma$ rays+0.02% SA | 2751.00 | 6.00 | - | 19.00 | 13.00 | 12.00 | 10.00 | 60.00 | 2.18 | 0.80 |
| 300Gy $\gamma$ rays+0.03% SA | 2649.00 | - | 16.00 | 9.00 | - | - | 4.00 | 29.00 | 1.09 | 0.37 |
| 400Gy $\gamma$ rays+0.04% SA | 2500.00 | - | 17.00 | 7.00 | - | 14.00 | 0.00 | 38.00 | 1.52 | 0.52 |
| **Total** | **10720.00** | **12.00** | **33.00** | **35.00** | **28.00** | **26.00** | **23.00** | **157.00** | **5.86** | **2.13** |

**Table 10.** Frequency and spectrum of morphological mutants

| Treatments | N | P | G | L | F | P | S | TMP | Mutated plants | k |
| --- | --- | --- | --- | --- | --- | --- | --- | --- | --- | --- |
| Control | 2960.00 | - | - | - | - | - | - | - | - | - |
| 100Gy $\gamma$ rays | 2910.00 | 10.00 | - | 8.00 | 2.00 | - | 9.00 | 29.00 | 1.00 | - |
| 200Gy $\gamma$ rays | 2851.00 | 2.00 | 9.00 | 14.00 | 4.00 | 11.00 | 10.00 | 50.00 | 1.75 | - |
| 300Gy $\gamma$ rays | 2807.00 | - | 14.00 | 8.00 | 5.00 | - | 5.00 | 32.00 | 1.14 | - |
| 400Gy $\gamma$ rays | 2755.00 | 7.00 | 10.00 | - | 4.00 | 8.00 | 3.00 | 32.00 | 1.16 | - |
| **TOTAL** | **11323.00** | **19.00** | **33.00** | **30.00** | **15.00** | **19.00** | **27.00** | **143.00** | **5.05** | - |
| 0.01%SA | 2890.00 | 10.00 | - | 11.00 | 12.00 | - | 4.00 | 37.00 | 1.28 | - |
| 0.02%SA | 2807.00 | 7.00 | 13.00 | 6.00 | 5.00 | 9.00 | 8.00 | 48.00 | 1.71 | - |
| 0.03%SA | 2757.00 | - | - | 8.00 | 10.00 | 9.00 | 9.00 | 36.00 | 1.31 | - |
| 0.04%SA | 2630.00 | 11.00 | 8.00 | 10.00 | 10.00 | - | 7.00 | 46.00 | 1.75 | - |
| **TOTAL** | **11084.00** | **28.00** | **21.00** | **35.00** | **37.00** | **18.00** | **28.00** | **167.00** | **6.05** | - |
| 100Gy $\gamma$ rays +0.01% SA | 2730.00 | 8.00 | - | - | 14.00 | - | 10.00 | 32.00 | 1.17 | 0.46 |
| 200Gy $\gamma$ rays+0.02% SA | 2661.00 | 8.00 | - | 17.00 | 15.00 | 13.00 | 11.00 | 64.00 | 2.41 | 0.60 |
| 300Gy $\gamma$ rays+0.03% SA | 2559.00 | 3.00 | 15.00 | 11.00 | - | 2.00 | 5.00 | 36.00 | 1.41 | 0.48 |
| 400Gy $\gamma$ rays+0.04% SA | 2410.00 | - | 14.00 | 9.00 | - | 15.00 | 0.00 | 38.00 | 1.58 | 0.50 |
| **TOTAL** | **10360.00** | **19.00** | **29.00** | **37.00** | **29.00** | **30.00** | **26.00** | **170.00** | **6.56** | **2.04** |

N = Number of M<sub>2</sub> plants, TMP = Total mutated plants, k = coefficient of interaction based on % M<sub>2</sub> mutated plants.

**Table 11.**
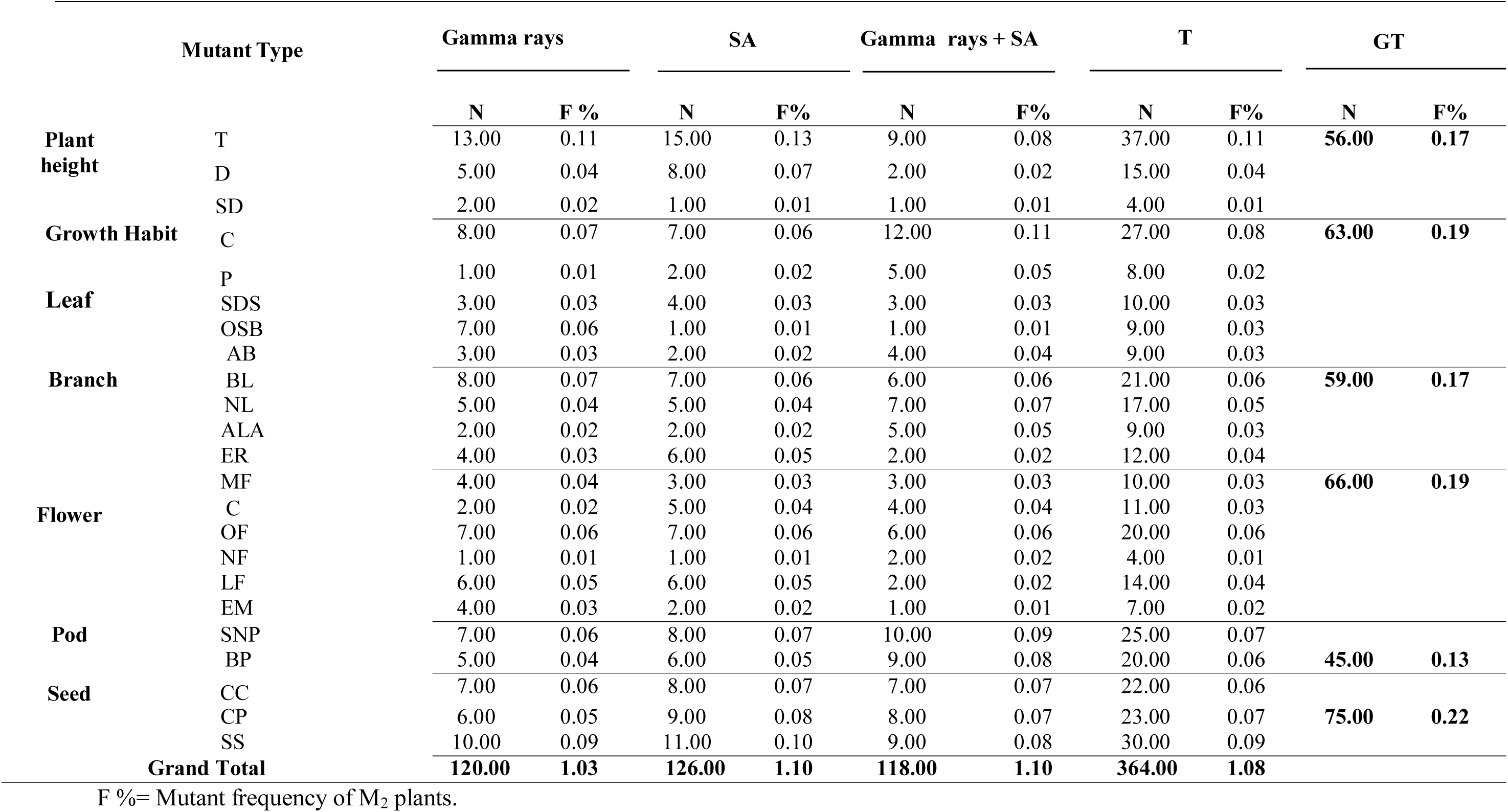
Frequency and spectrum of morphological mutants induced by various mutagens in M_2_ Gomati VU-89.

**Table 12.** Frequency and spectrum of morphological mutants induced by various mutagens in M_2_ generation of cowpea var. Pusa-578.

| Mutant Type |  | Gamma rays |  | SA |  | Gamma rays + SA |  | Total |  | GT |  |
| --- | --- | --- | --- | --- | --- | --- | --- | --- | --- | --- | --- |
|  |  | N | F % | N | F% | N | F% | N | F% | N | F% |
| **Plant height** | T | 10.00 | 0.09 | 11.00 | 0.10 | 8.00 | 0.08 | 29.00 | 0.09 | **44.00** | **0.13** |
|  | D | 4.00 | 0.04 | 7.00 | 0.06 | 1.00 | 0.01 | 12.00 | 0.04 |  |  |
|  | S | 1.00 | 0.01 | 1.00 | 0.01 | 1.00 | 0.01 | 3.00 | 0.01 |  |  |
| **Growth habit** | C | 7.00 | 0.06 | 6.00 | 0.05 | 11.00 | 0.11 | 24.00 | 0.07 | **57.00** | **0.17** |
|  | P | 4.00 | 0.04 | 1.00 | 0.01 | 6.00 | 0.06 | 11.00 | 0.03 |  |  |
|  | SDS | 2.00 | 0.02 | 3.00 | 0.03 | 2.00 | 0.02 | 7.00 | 0.02 |  |  |
|  | OSB | 5.00 | 0.04 | 2.00 | 0.02 | 1.00 | 0.01 | 8.00 | 0.02 |  |  |
|  | AB | 2.00 | 0.02 | 3.00 | 0.03 | 2.00 | 0.02 | 7.00 | 0.02 |  |  |
| **Leaf** | BL | 7.00 | 0.06 | 6.00 | 0.05 | 5.00 | 0.05 | 18.00 | 0.05 | **50.00** | **0.15** |
|  | NL | 4.00 | 0.04 | 4.00 | 0.04 | 5.00 | 0.05 | 13.00 | 0.04 |  |  |
|  | ALA | 1.00 | 0.01 | 3.00 | 0.03 | 4.00 | 0.04 | 8.00 | 0.02 |  |  |
|  | ER | 5.00 | 0.04 | 5.00 | 0.05 | 1.00 | 0.01 | 11.00 | 0.03 |  |  |
| **Flower** | MF | 3.00 | 0.03 | 2.00 | 0.02 | 4.00 | 0.04 | 9.00 | 0.03 | **64.00** | **0.20** |
|  | C | 3.00 | 0.03 | 4.00 | 0.04 | 5.00 | 0.05 | 12.00 | 0.04 |  |  |
|  | OF | 6.00 | 0.05 | 8.00 | 0.07 | 5.00 | 0.05 | 19.00 | 0.06 |  |  |
|  | NF | 2.00 | 0.02 | 2.00 | 0.02 | 1.00 | 0.01 | 5.00 | 0.02 |  |  |
|  | LF | 5.00 | 0.04 | 5.00 | 0.05 | 1.00 | 0.01 | 11.00 | 0.03 |  |  |
|  | EM | 3.00 | 0.03 | 4.00 | 0.04 | 1.00 | 0.01 | 8.00 | 0.02 |  |  |
| **Pod** | SNP | 5.00 | 0.04 | 7.00 | 0.06 | 9.00 | 0.09 | 21.00 | 0.06 | **45.00** | **0.14** |
|  | BP | 4.00 | 0.04 | 5.00 | 0.05 | 8.00 | 0.08 | 17.00 | 0.05 |  |  |
| **Seed** | CC | 6.00 | 0.05 | 7.00 | 0.06 | 6.00 | 0.06 | 19.00 | 0.06 | **75.00** | **0.23** |
|  | CP | 5.00 | 0.04 | 8.00 | 0.07 | 7.00 | 0.07 | 20.00 | 0.06 |  |  |
|  | SS | 9.00 | 0.08 | 10.00 | 0.09 | 8.00 | 0.08 | 27.00 | 0.08 |  |  |
| **Grand Total** |  | **103.00** | **0.91** | **114.00** | **1.03** | **102.00** | **0.98** | **319.00** | **0.97** |  |  |

F% = Mutant frequency of M<sub>2</sub> Plants

#### I. Plant height mutants (**Fig 5**)

i. Tall mutants: Such mutants were induced with frequency of 0.11 and 0.09% at lower doses of gamma rays and SA in both the cowpea varieties. The mutants Plants attained a height 180-185 cm compared to 178-183 cm in control plants.
ii. Dwarf mutants: Such mutants were mostly induced with a 0.04% frequency of the total morphological mutations. These mutants were induced at higher doses of all the treatments except the combination treatments in the var. Gomati VU-89. These mutants showed lesser leaves, reduced pod and seed size and poor yield. The plant height was severely reduced and it ranged from 110-115 cm.
iii. Semi dwarf mutants: Such mutants were induced at 0.01% frequency. Pod length was increased, however, the seeds were smaller in size.

#### II. Growth habit mutants (Fig 5)

i. Compact/Bushy mutants: Such mutants are characterised with 150.55 cm average height, robust growth. The yield per plant was slightly higher and matured five days earlier compared to control plants. Such mutants were mostly induced by the moderate and higher treatments. The bushy mutants appeared at 0.08% of the total morphological mutations in the var. Gomati VU-89 and 0.07% in var. Pusa-578.
ii. Prostrate mutants: Such mutants were flat on ground, branches spread, possess long internodes, and reflected trailing tendency at the soil surface. They possess, less number of pods containing 6-7 shriveled seeds with rough and hard seed coat. In both the varieties of cowpea prostrate mutants were commonly induced with combination treatments
iii. Spreading mutants: Such mutants were observed in 200Gy gamma rays treatment (Gomati VU-89) and 0.03% SA treatment (Pusa-578). The leaves were mostly broad and branches appeared spreading. Seed yield per plant was significantly reduced. Their frequency was 0.03% and 0.02% in varieties Gomati VU-89 and Pusa-578, respectively.
iv. One sided branching mutants: Mutant plants had thick stems bearing branches on one side and were late in flowering and maturing, low yielded with few pods and shriveled seeds. These mutants were screened in SA doses in both the cowpea varieties and in combination treatment of 200Gy gamma rays + 0.02% SA in the var. Pusa-578.

#### III. Leaf mutants (Fig 6-7)

i. Broad/Giant leaves mutants: Moderate doses of gamma rays and SA treatments induced higher frequency of broad leaf mutants.
ii. Narrow leaves mutants: Such mutants possess narrow leaves with pointed leaf tips. Compared to control, the yield was lower, small pods with few seeds per pod. Higher doses of mutagens induce higher frequency of mutants in both the cowpea varieties.
iii. Altered leaf architecture mutants: Plants showed reduced height (about 95 cm) at maturity. Such mutants were induced by all the mutagen treatments
iv. Elongated rachis: Mutants possessed increased length of rachis with narrow leaflets.

They appeared mostly at the lower mutagen doses with a frequency of 0.04% and 0.03% in the var. Gomati VU-89 and var. Pusa-578, respectively.

**Fig 5.**
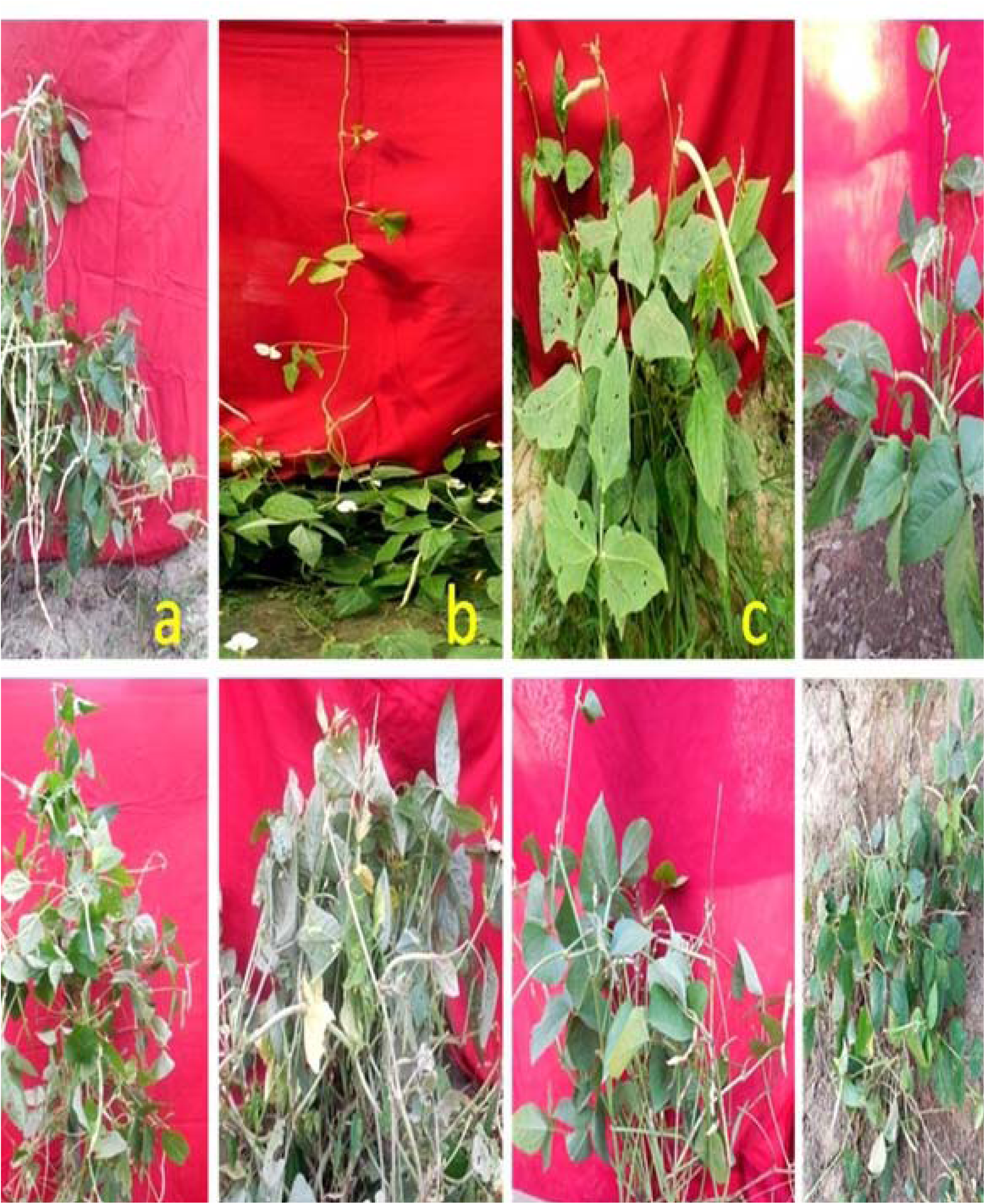
**Morphological mutations.** **a** Control Plant **b** Tall mutant **c** Dwarf mutant **d** Semi dwarf mutant **e** Spreading mutant. **f** Bushy mutant. **g** Axillary branched mutant. **h** Prostrate mutant

**Fig 6.**
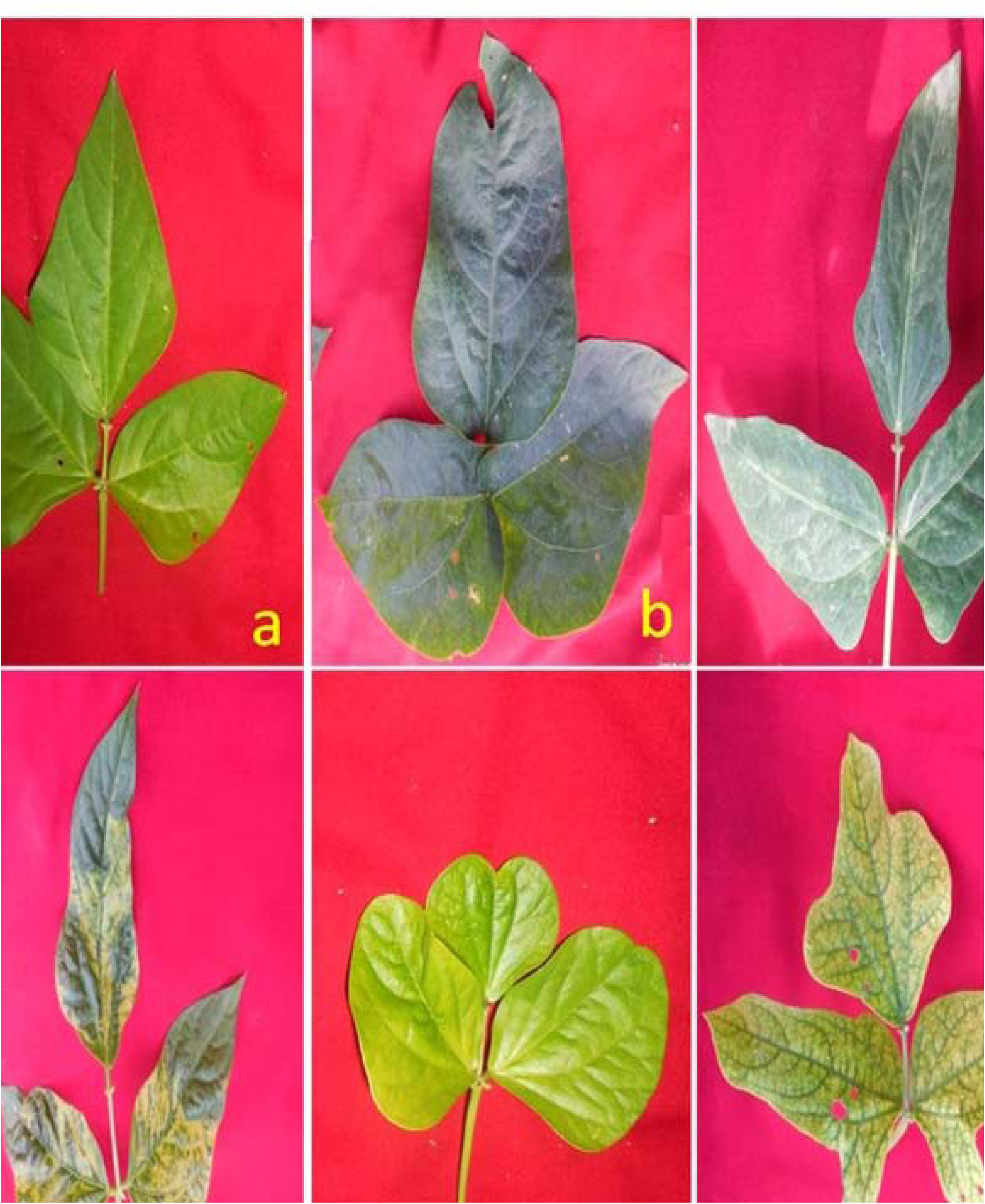
**Morphological mutations in leaves** **a** Control leaf **b** Broad leaf mutant **c** Narrow leaf mutant **d** Long rachis mutant **e** Trifoliate mutant **f L**eaf architecture.

**Fig 7.**
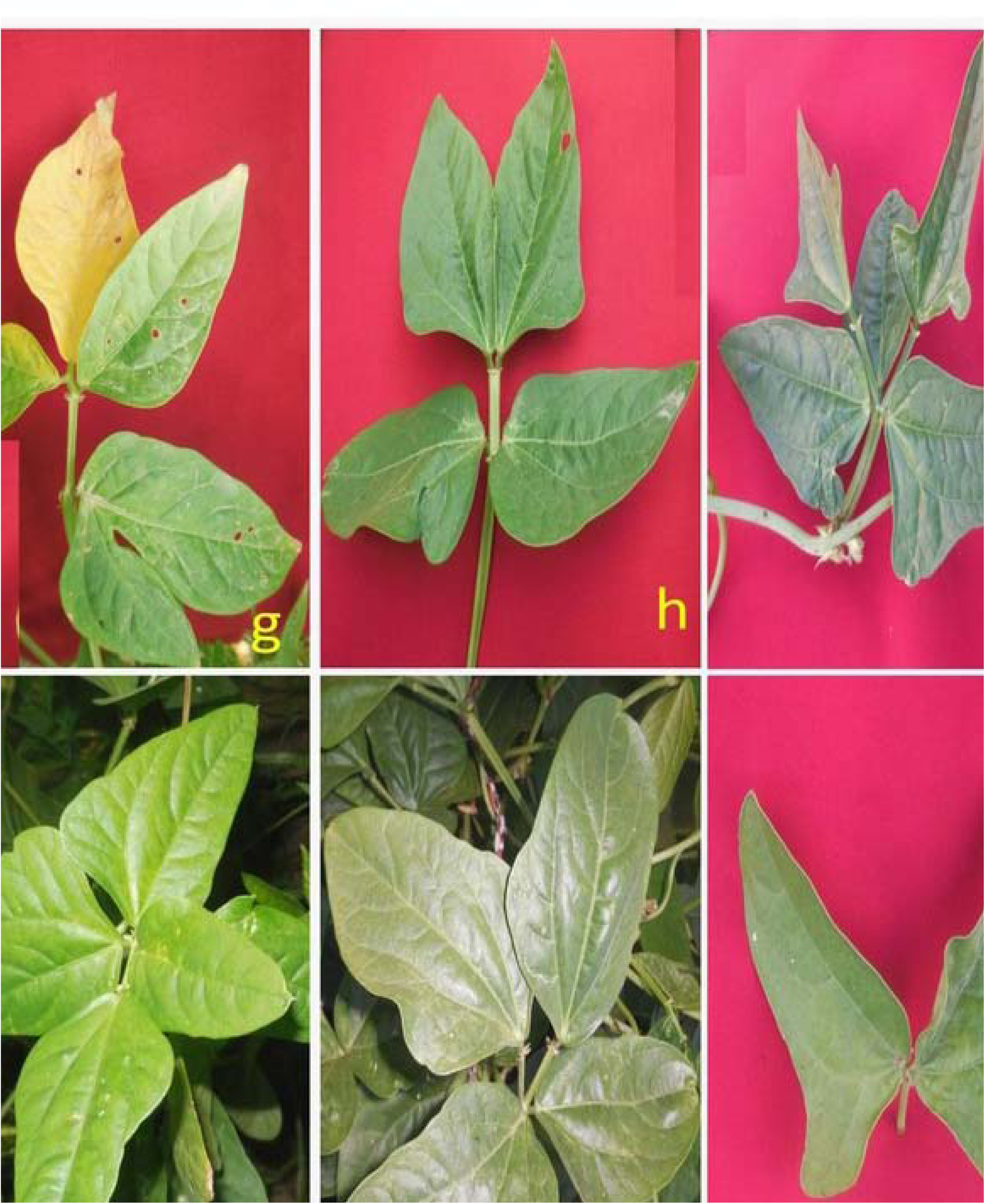
**Morphological mutations (plants with altered leaflet number)** **g** Mutant showing incision of leaf. **h** Tetrafoliate with fused terminal leaflets. **i** Pentafoliate with variations in leaflet size and shape. **j** Tetrafoliate with free leaflets. **k** Broad tetrafoliate leaflets. **l** Bifoliate leaf.

#### IV. Flower mutants (Fig 8)

i. Multiple flower mutants: Such mutants have small pods and shrivelled seeds. Mutants were late in maturity compared to the control. These mutants appeared at frequency of about 0.03% in both the varieties.
ii. Flower colour mutants: The 300 Gy gamma rays showed higher frequency of white or blue flower colour mutants. Their frequency was recorded as 0.03% in the var. Gomati VU-89 and 0.04% in the var. Pusa-578.
iii. Open flower mutants: Such mutants app ea red at 0.06% frequency of total mutation at moderate and higher doses of SA in var. Gomati VU-89 and Pusa-578.
iv. Non-flowering/Vegetative mutants: These mutants were noticed in SA and gamma rays+SA treatments.
v. Late flowering mutants: Higher doses of mutagens induced large number of late flowering mutants.
vi. Early maturity mutants: Such mutants matured 10-15 days earlier than control and were induced at the lower mutagens doses used singly or in combination. The frequency of such mutants was 0.02% in both the varieties.

#### V. Pod mutants (Fig 9)

i. Small/Narrow pods: Such mutants possessed small and /or narrow pods compared to long pods in control, observed in combined mutagen treatments. The seed yield per plant was also reduced as compared to control plants.
ii. Bold seeded pod mutants: The bold size of pods led to significant increase in yield per plant as compared to control. Lower and moderate doses of individual and combined mutagens induced broad spectrum of these mutants.

#### VI. Seed mutants (Fig 10)

i. Seed coat colour mutants: Such mutants are characterized red and black seed coats compared to brown and white seed coats in control plants of the var. Gomati VU-89 and var. Pusa-578, respectively.
ii. Seed shape and surface mutants: Seeds of control plants of both the varieties were of kidney shaped with rough seed surface and were induced in lower doses. Mutants with globular shape and smooth seed surface were isolated in the mutagen treated population.

## 4. Discussion

A considerable decline in seed germination, seedling height, pollen fertility and plant survival of treated population was observed in mutagen treatments. However, the extent of diminution in the bio-physiological parameters differed among mutagen doses. The diminution in plant survival was recorded in all the mutagen doses, and confirmed the fact that the mutagens interact independently with the genes governing a particular trait and that mutations occur in a random manner. These results are in line with earlier reports on several grain legumes [15], [16], [17], [18], [19], [20], [21]. However, the promoting effects of mutagen doses on bio-physiological parameters were also reported in several other crops [22], [23], [24], [25], [26].

**Fig 8.**
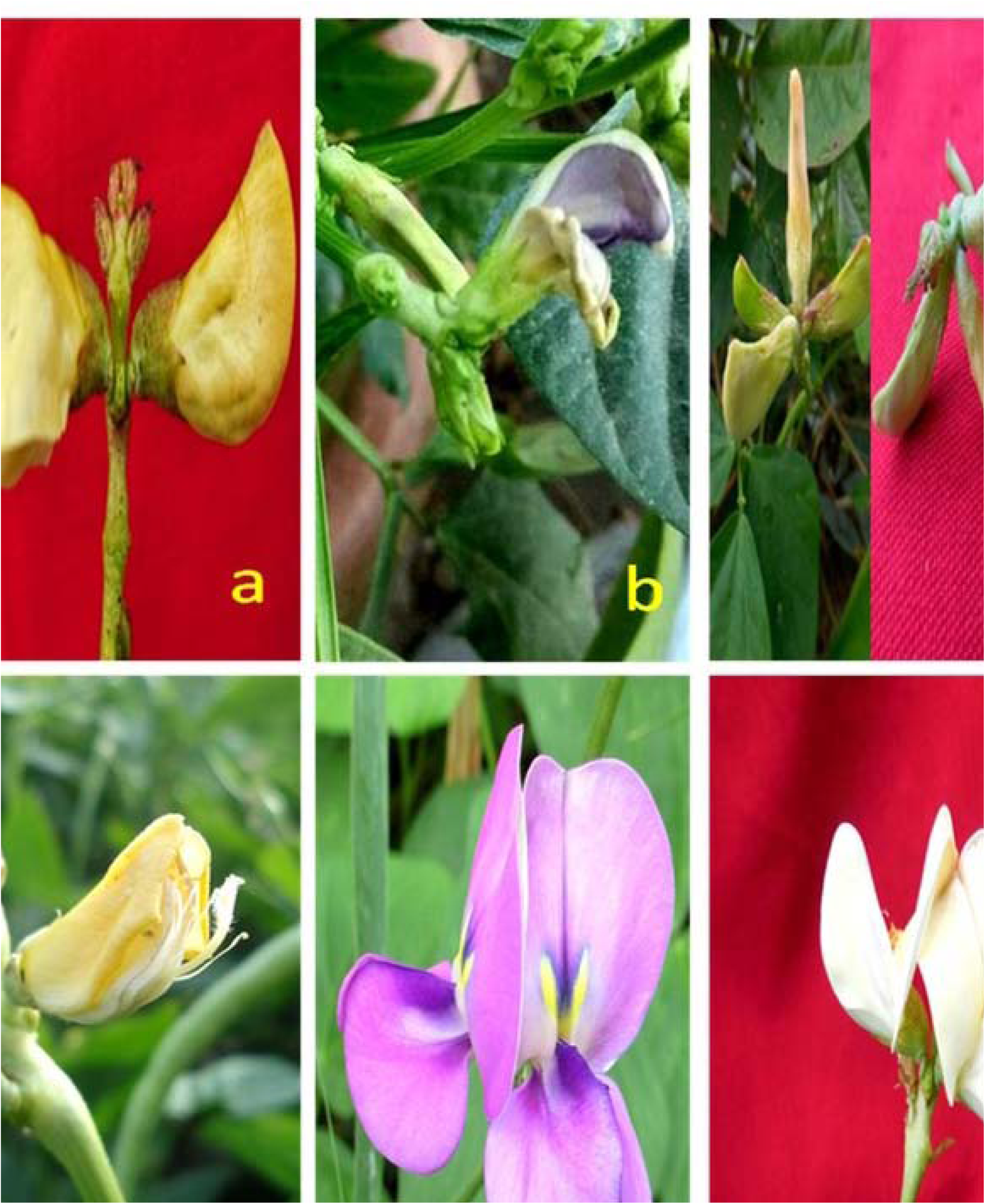
**Morphological mutations in flowers** **a** Control flowers **b P**etal mutants **c** Multiple flowers. **d** Open flower mutant. **e** Light blue colour flower mutant. **f** White flowers mutant.

**Fig 9.**
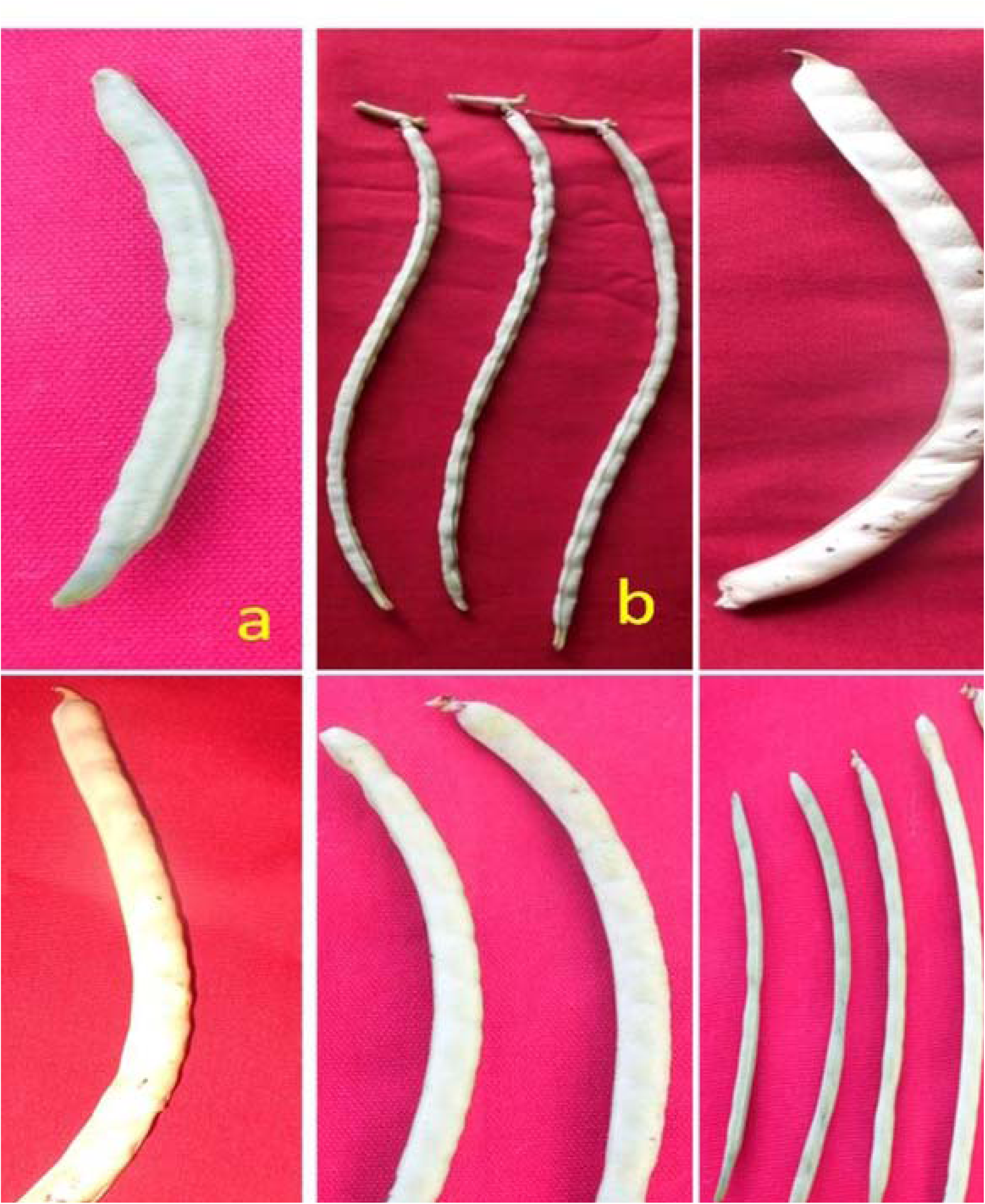
**Morphological mutations in pods** a Normal pod (control). b Long pods. c Broad pod with bold seeds. d Broad pods. e Pod width mutants. f Pod length mutants.

**Fig 10.**
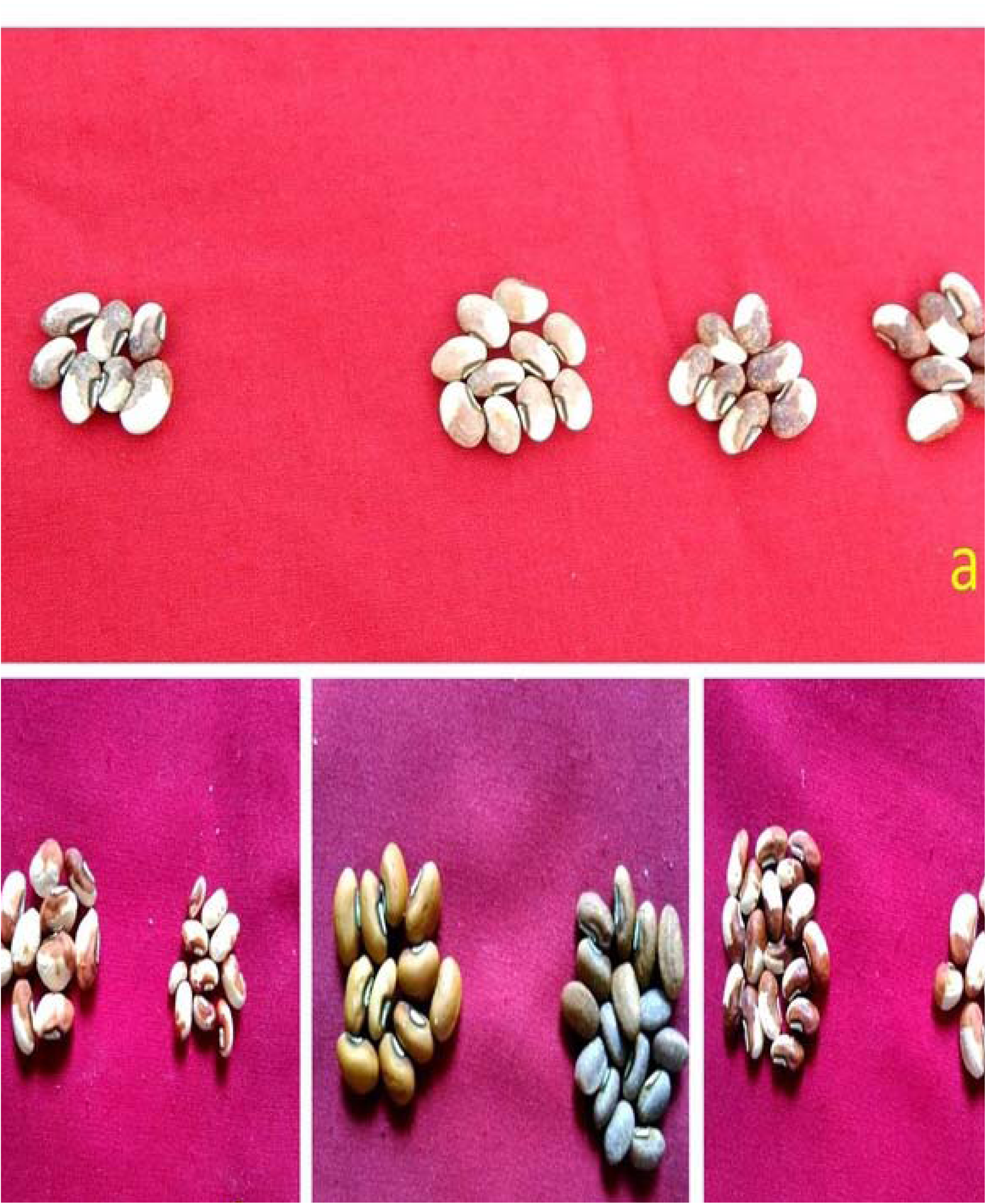
**Morphological mutations in seed characters)** **a S**eed coat mutants. **b** Bold seed mutants **c** Red seed mutants. **d** Wrinkled seed mutants.

### 5.1. Seed Germination

The assessment of seed germination gives an idea about the mutagenic potency, the dose at which germination is below 50 per cent is treated as lethal or undesirable. Since, the purpose of this present study was to develop large mutagenized population to facilitate effective screening of the high yielding mutants, therefore, the lethal doses of mutagens were discarded to ensure high frequency of desirable mutation. Decreased germination due to mutagen treatments may be the attributed to mutagen induced germination inhibition. The inhibition in seed germination was found to be more or less comparable in gamma rays and SA treatments, while the combined treatments induced higher seed germination inhibition in both the cowpea varieties, which may be attributed to the synergistic effect of the two mutagens. Chauhan and Singh [27] reported that gamma rays interact and cause disruption of tunica zone that leads to the decrease in seed germination. Alterations in hormone synthesizing enzymes may be another reason for seed germination inhibition [28]. In the present study, the decrease in pollen fertility has been noticed in the mutagenized population. Pollen sterility was maximum in combined mutagen doses than individual mutagen doses. This may cause meiotic anomalies and leading to the formation of aberrant pollen grains. Reduction in pollen fertility after mutagen treatments have also been reported in lentil [29], [30], cowpea [31], fenugreek [32], faba bean [33]. The pollen sterility percentage was found remarkably less in M_2_ than M_1_ generation. The survival at maturity was declined with increasing mutagen dose in both the varieties of cowpea. These findings support the earlier reports in *Lathyrus sativus* [34] in *Capsicum annum*, [35]. The reduction of plant survival at maturity may be attributed to the disturbed physiological processes [36], [37] or cytological damages Manjaya [38]. The observations recorded on several bio-physiological and cytological parameters revealed that cowpea var. Pusa-578 was comparatively more sensitive towards the individual and combination doses than Gomati VU-89. The differential response of varieties from the same species towards the mutagens have also been reported earlier in *Cajanas cajan* [39], *Vigna* spp [40], [41], [42], *Phaseolus vulgaris* [43], [44], *Jatropha curcus* [45] and *Lens culinaris* [46]. Sparrow *et al.* [47] suggested that differential genotypic sensitivity towards mutagens may be due to the gene loci mutated by one mutagen in one variety were not necessarily mutated by the second mutagen in another variety. The two cowpea varieties viz., Gomati VU-89 and Pusa-578, used in the present study, were from different races, therefore, it is obvious that the two varieties have substantial variation in their genome and thereby the heritable genetic identity of the two varieties is immensely different. Although the spectrum of variations was more or less similar in both the varieties but the mutagen doses to achieve that were not identical in various parameters studied. It can be concluded from the present bio-physiological that mutagenesis induced a comparable spectrum of apparent variation through random mutation in both the varieties of diverse origin with the variable treatments of gamma rays and SA.

### 5.2. Chlorophyll mutations

In mutation breeding, the increase in mutation frequency is necessary for crop improvement. In order to acquire desired mutations at highest rates, the picking of optimum mutagen and appropriate dose is important. Both the varieties of cowpea showed a differential response towards gamma rays, SA and gamma rays+SA treatments. The chlorophyll mutants were identified as per the categorization of Gustafsson [9]. Gamma rays+SA caused maximum frequency of chlorophyll mutations than the gamma rays and SA. Similar trends were reported in chickpea [48], *Oryza sativa* [49], *Vigna mungo* [50] *Zingiber officinale* [51], *Lablab purpureus* [52]. On contrary Pavadai *et al*. [53] in soybean, Barshile *et al*. [54] in chickpea, Kumar *et al*. [55] in mungbean reported the linear increase in chlorophyll mutation frequency with increase in doses of different mutagen treatments. In the present study, albina and chlorina mutants were found more frequently in both the cowpea varieties. SA was found to be more potent in the production of albina and chlorina mutants than gamma rays. The gamma rays+SA treatments have induced maximum frequency of albina and chlorine mutants than gamma rays and SA treatments employed individually. The synergistic effects in the combination treatments are very much evident from the higher frequency of chlorophyll mutations. The possible reason for synergism might be that the mutagen first applied may expose the accessible protected mutable sites to the second mutagen, the repair enzymes may be rendered non functional by the second mutagen which indirectly facilitates mutations fixation [11]. The synergism were earlier documented in *Cicer arietinum* [56], *Phaseolus vulgaris* [57], *Vigna spp.*, [58], *Pisum sativum* [59], *Linum usitatissimum* [60].

### 5.3. Mutagenic effectiveness and efficiency

Induced mutagenesis has been known as a vital tool for the improvement of wide range of crops. The success of mutation breeding depends on enhancing the efficiency of desired mutation. Further the mutagen choice is imperative to achieve the desired mutation rate [12], [61]. Whereas mutagenic efficiency on the contrary the SA doses were superior among mutagen doses employed in present study. Similar results have been reported by Balai and Krishna [62] while studying the effectiveness of ethylmethane sulphonate, sodium azide and hydroxylamine on mungbean. Earlier workers such as Kaul and Bhan [63] in rice, Laskar and Khan [64] in lentil, Wani [65] and Shah *et al*. [66] in chickpea, Khan and Tyagi [67] in soybean and Dhulgande *et al*. [68] in pea have also reported the more effectiveness of chemical mutagens over physical mutagens [69] while studying the mutation breeding in cowpea. As per them, in order to attain higher effectiveness and efficiency, the mutation effect was more prominent in the cell that led to meiotic anomalies. Our results reflected a differential effectiveness in two varieties of cowpea which depict the genetic divergence and mutagenic sensitivity towards different mutagen doses. In the present study, the gamma rays+SA were more efficient than individual gamma rays and SA treatments andthis may be attributed to the higher chlorophyll mutants. On contrary Zeerak [70] in brinjal reported that gamma rays+ EMS doses were less efficient than the individual doses of gamma rays and EMS. In the present study, lower and medium doses of gamma rays and SA and gamma rays+SA treatments were found to be more efficient in both the varieties of cowpea. The results were in line with the earlier results in *Lens culinaris* [64] *Glycine max* [67], *Vigna mungo* [71]. However, contrary results of linear increase in mutagenic efficiency with the increase in the mutagen concentrations have also been reported in black gram [72] and millet [73]. Makeen *et al*. [74] also reported higher 613 mutagenic efficiency at lower and medium doses in urdbean.

### 5.4. Morphological mutations

Even though the majority of the induced morphological mutants were uneconomical, however, these mutants can serve as a source of valuable genes in hybridisation programmes. Toker [75] reported that the morphological mutants are useful in gene mapping and phylogentic studies of crops. Mutagen induced chromosomal anomalies or the pleiotropic effects of mutated genes could be the probable reason for the induction of morphological mutants [76], [77]. Different mutagen doses and duration of treatment may be attributed to the relative proportion of different mutation types [78].. Khursheed *et al*. [79] reported maximum morphological mutants frequency in gamma rays and lowest with combined treatments while investigating the frequency and spectrum of M_2_ morphological faba bean mutants. Previous reports of a high frequency and boarder spectrum of morphological mutations in crops such as *Vigna mungo* [41], *Cicer arietinum* [80], [81], *Lens culinaris* [82], *Phaseolus vulgaris* [83] *Glycine max* [67] also confirm that mutagens are effective in inducing morphological mutations in wide range of crops that may serve as a source of valuable genes for the crop improvement programs. Some morphological mutants (early maturing, dwarf, bushy, bold seeded pods and639 flower colour), screened in M_2_ generation in the present study, can serve as source of 640 valuable genes or used as a parents in cross breeding programmes. Dwarf mutants, as 641 recorded in the present study, have been reported earlier in chickpea [21] barley [84], 642 mungbean spp. [77], lentil [85]. The decreased length of internodes and/or number of 643 internode could be attributed to the induction of dwarf and bushy mutants [86]. In our 644 experimentation dwarf and bushy mutants showed substantial decrease in internode length 645 but number of internodes were similar to that of control plants. The mutated genes that 646 control the altered plant height and growth habits are monogenic recessives [87]. Whereas polygenes govern the semi dwarf trait in mutants in wheat [12] and triticale [88]. However, Qin *et al*. [89] reported a dominant mutation in a single gene in dwarf mutants in rice. Both dwarfness and bushy habit of a genotype contribute to the production and productivity. The bold pod mutants are associated with high yield which may be valuable in increasing seed size that lead to enhanced cowpea yield. Singh [90] reported that gene mutations govern the bold seeded trait in *Vigna mungo*.

## Conclusions

Increase in mutagen doses lead to a progressive decline in the values of all bio-physiological parameters. Combination treatments showed an enhanced effect on seed germination, seedling height and pollen fertility compared to individual mutagen treatments. A dose dependant increase in the frequency of chlorophyll mutations was observed in both the varieties. The estimation of coefficient of interaction (k) did not show any additive effects of mutagens. Mutagenic effectiveness was highest at the moderate doses of gamma rays and SA, however, lower doses of gamma rays+SA were most effective and efficient in both the varieties. The effectiveness of mutagens was found as SA> gamma rays + SA> gamma rays. Mp/I and Mp/S based mutagenic efficiency was gamma rays> gamma rays+ SA> SA whereas Mp/Me based order was gamma rays + SA> gamma rays> SA. A diverse range of mutagen induced morphological mutants were obtained in the population of two varieties of cowpea in M_2_ generation, many of which are useful from yield improvement prospective. Of all the mutant types, seed and flower mutants followed by growth habit, plant height and pod length were of maximum occurrence in the two varieties. The spectrum of morphological mutations induced by combined mutagens was relatively boarder than gamma rays and SA treatments in both the varieties.

## Author Contributions

**Conceptualization:** Aamir Raina, Samiullah Khan.

**Data curation:** Aamir Raina.

**Formal analysis:** Aamir Raina.

**Investigation:** Aamir Raina.

**Methodology:** Aamir Raina.

**Project administration:** Aamir Raina.

**Resources:** Aamir Raina.

**Software:** Aamir Raina.

**Supervision:** Samiullah Khan.

**Validation:** Aamir Raina.

**Visualization:** Aamir Raina.

**Writing - original draft:** Aamir Raina.

**Writing - review & editing:** Aamir Raina.

